# The efficacy of a ketogenic diet in preclinical models of IDH wild-type and IDH mutant glioma

**DOI:** 10.1101/2021.09.09.459672

**Authors:** Rodrigo Javier, Wenxia Wang, Michael Drumm, Kathleen McCortney, Jann N. Sarkaria, Craig Horbinski

## Abstract

Infiltrative gliomas are the most common neoplasms arising in the brain, and remain largely incurable despite decades of research. A subset of these gliomas contains mutations in *isocitrate dehydrogenase 1* (*IDH1*) or, less commonly, *IDH2* (together called “IDH^mut^”). These mutations alter cellular biochemistry, and IDH^mut^ gliomas are generally less aggressive than IDH wild-type (IDH^wt^) gliomas. Some preclinical studies and clinical trials have suggested that a ketogenic diet (KD), characterized by low-carbohydrate and high-fat content, may be beneficial in slowing glioma progression. However, not all studies have shown promising results, and to date, no study has addressed whether IDH^mut^ gliomas might be more sensitive to KD. The aim of the current study was to compare the effects of KD in preclinical models of IDH^wt^ versus IDH^mut^ gliomas. *In vitro*, simulating KD by treatment with the ketone body β-hydroxybutyrate had no effect on the proliferation of patient-derived IDH^wt^ or IDH^mut^ glioma cells. Likewise, a cycling KD, wherein mice alternated between KD and a standard diet (SD), had no effect on the *in vivo* growth of patient-derived IDH^wt^ or IDH^mut^ gliomas, even though the cycling KD did result in persistently elevated circulating ketones. Furthermore, KD conferred no survival benefit in mice engrafted with Sleeping-Beauty transposase-engineered IDH^mut^ glioma. These data suggest that neither IDH^wt^ nor IDH^mut^ gliomas are particularly responsive to KD.

## INTRODUCTION

Diffusely infiltrative gliomas strike over 17,000 people in the United States per year [1]. The vast majority of these tumors recur and progress, despite advancements in surgery, chemotherapy, radiotherapy, and immunotherapy. In 20-39 year-olds, gliomas are the 2^nd^ most common cause of cancer death in men, and are the 5^th^ most common cause in women [1]. The most common type of primary brain cancer in adults is diffusely infiltrative glioma, and the most common subtype of infiltrative glioma, glioblastoma (GBM), is unfortunately also the most lethal. Despite great advances in treating many other kinds of cancer, the median survival of GBM patients is still only 12-15 months after diagnosis, even with surgical resection, radiation, and temozolomide therapy. Long-term prognosis is grim; only about 15% of patients with an infiltrative glioma survive 5 years after diagnosis [2, 3]. As a group, primary brain cancers rank #1 among all cancers in terms of average years of life lost [4]. Despite intensive research, only modest progress has been made in the treatment of gliomas, and new approaches are badly needed.

Alterations in cell metabolism have long been known to be a hallmark of cancer, ever since Otto Warburg first described the preferential reliance on aerobic glycolysis over oxidative phosphorylation in cancer cells [5, 6]. However, it was only relatively recently that mutations in metabolic genes were found to occur in some cancers. For example, approximately 20-30% of infiltrative gliomas carry mutations in isocitrate dehydrogenase 1 (IDH1) or, far less commonly, IDH2 (together referred to as “IDH^mut^”) [7-9]. This subset of gliomas tends to occur in grade 2-3 gliomas, disproportionately arises in younger adults, and is associated with longer survival. Wild-type IDH1 and IDH2 encode enzymes that catalyze the oxidative decarboxylation of isocitrate to α-ketoglutarate in the cytosol/peroxisomes and mitochondrion, respectively. In the process, these enzymes also generate reduced nicotinamide adenine dinucleotide phosphate (NADPH). Point mutations in codon 132 of IDH1 (usually R132H), and codon 172 of IDH2, cause the mutant enzymes to reduce α-ketoglutarate to D-2-hydroxyglutarate (D2HG), thereby consuming NADPH [10].

Because cancers mostly rely on glucose for energy and anabolism, numerous studies have explored the therapeutic potential of dietary carbohydrate restriction in cancer patients. This is achieved through a ketogenic diet (KD), which is characterized by high-fat, low-carbohydrate, and moderate-protein content. KD limits the bioavailability of carbohydrates and induces the liver to produce ketone bodies, such as β-hydroxybutyrate and acetoacetate, which are then converted into acetyl-CoA for use in the tri-carboxylic acid (TCA) cycle [11]. The goal of KD is to force tumor cells to use ketone bodies for energy, while still meeting the patient’s basic nutritional needs.

KD has been tested as an adjuvant therapeutic strategy in a number of cancers, including GBM, with mixed results thus far [12, 13]. However, an aspect of this research that has not yet been experimentally addressed is whether IDH^mut^ gliomas might be particularly responsive to KD. One study suggested that the D2HG product of IDH^mut^ can actually bind and inhibit ATP synthase, thereby inhibiting oxidative phosphorylation and ATP production [14]. In that study, human colorectal HCT116 cells transduced with IDH1 R132H were highly vulnerable to glucose deprivation *in vitro*, and were not able to use ketone bodies as effectively as IDH^wt^ HCT116 cells. A recent clinical study found that 8 weeks of KD did indeed elevate urinary ketones in glioma patients, with no significant change in fasting glucose or hemoglobin A1c. Additionally, the levels of intratumoral ketones were similar between IDH^wt^ and IDH^mut^ glioma patients [15]. However, that study was not designed to evaluate antitumor efficacy. Therefore, we sought to explore whether KD might preferentially inhibit the growth of IDH^mut^ gliomas *in vitro* and *in vivo*, using both patient-derived endogenous IDH1^wt^ and IDH1^mut^ xenografts in immunocompromised mice, as well as an isogenic Sleeping Beauty transposase-engineered model of IDH1^wt^ and IDH1^mut^ gliomas in immunocompetent mice.

## METHODS

### Ethics Statement

This study was performed in accordance with the recommendations in the Guide for the Care and Use of Laboratory Animals of the National Institute of Health. The protocol was approved by the Institutional Animal Care and Use Committee (IACUC) of Northwestern University (protocol #5715), and allowed for flank tumors to reach a volume of 6,000 mm^3^ (maximum dimensions of 20 mm × 30 mm). All surgery was performed under isoflurane inhalant anesthesia. Every effort was made to minimize animal suffering. Patient-derived cell lines were all developed under the auspices of an Institutional Review Board-approved protocol at Mayo Clinic and Duke University, with consent obtained from their donors.

### Cell Lines and Cell Cultures

Five cell types were derived from the Mayo Clinic Brain Tumor Patient-Derived Xenograft National Resource [16]. Three were IDH1^wt^ glioblastomas (GBM6, GBM12, and GBM43), and 2 were IDH1^mut^ grade 4 astrocytomas (GBM164 and GBM196). Two additional IDH1^mut^ cell types were TB09, a WHO grade 3 astrocytoma obtained from Dr Hai Yan at Duke University; and HT1080, a fibrosarcoma cell line from the American Type Culture Collection (the fibrosarcoma cell line was chosen to directly compare any possible differences in proliferation based solely on IDH1^mut^ status). All IDH1^mut^ cells were R132H except for HT1080, which was R132C IDH1. NPA and NPAC1 are rodent isogenic cell lines engineered using the Sleeping Beauty transposase system, courtesy of Dr Maria Castro from the University of Michigan [17]. Both NPA and NPAC1 have activating mutations in *NRAS* and inactivating mutations in *TP53* and *ATRX*; NPAC1 also expresses IDH1 R132H [17]. All IDH1^mut^ cell types produced high amounts of D2HG via liquid chromatography-mass spectrometry, relative to the IDH1^wt^ cells (not shown). All cell types were authenticated via short tandem repeat analysis, and are summarized in **Table 1**.

**Table 1:**
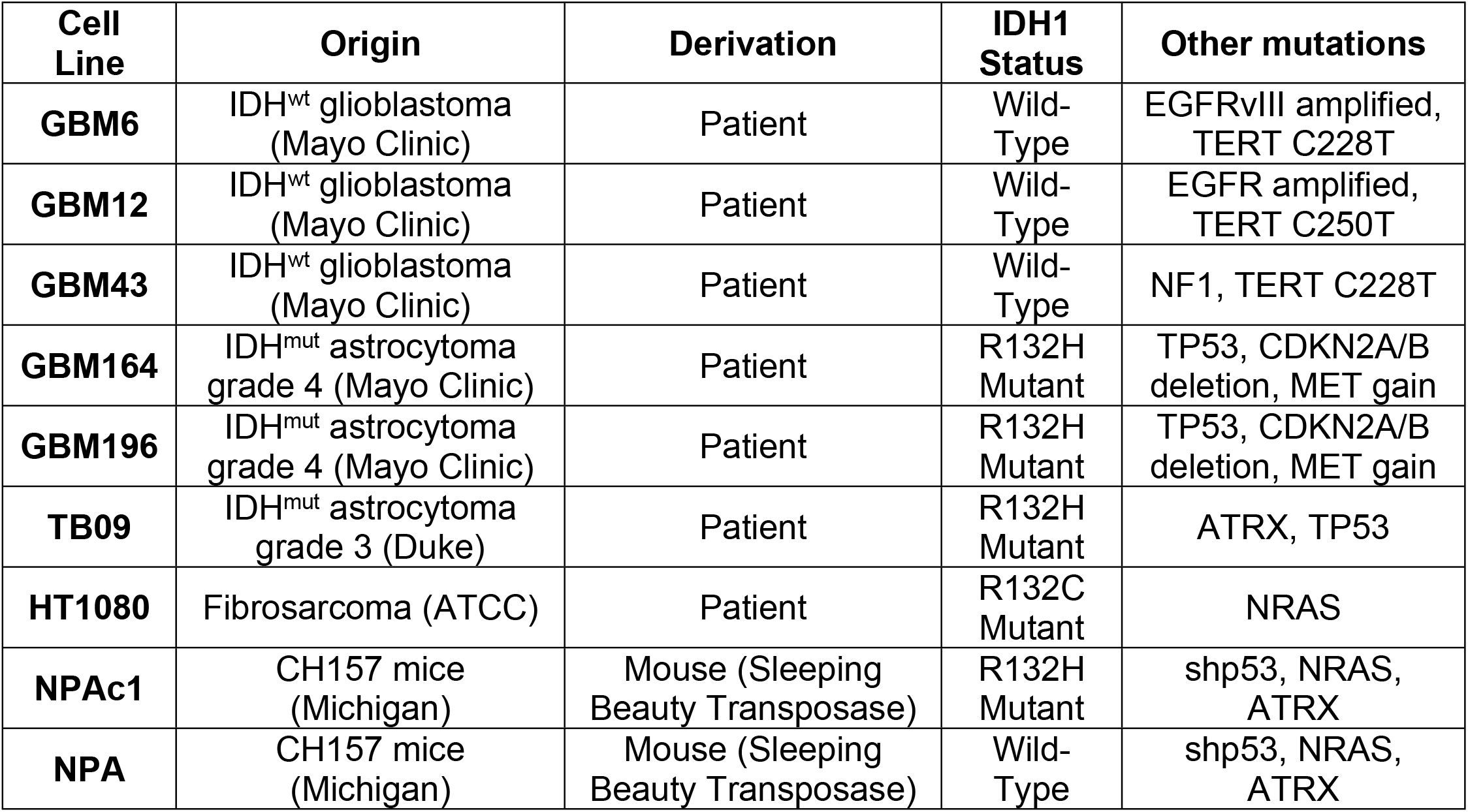
Summary of cell lines used.

For cell culture studies, GBM6, GBM43, TB09, and HT1080 cells were grown in Dulbecco’s modified eagle medium (DMEM, Corning) supplemented with 10% fetal bovine serum and 1% penicillin streptomycin at 37°C with 5% CO2.

### In vitro proliferation

Cells were plated in 24 well plates at a concentration of 5×10^4^ cells per well in triplicate. Experimental wells were supplemented with 10 mM β-hydroxybutyrate (Sigma Product #H6501). To mimic the low glucose environment characteristic of physiological ketosis, a formulation of DMEM (Corning) with 1.0 g/L of glucose was tested alongside the normal glucose concentration of 4.5 g/L. Plates were trypsinized at specific time points, and live cells were counted via trypan blue exclusion using a BioRad TC20 Automated Cell Counter. Absolute cell counts were used to determine cell viability rather than the 3-(4,5-dimethylthiazol-2-yl)-2,5-diphenyltetrazolium (MTT) assay, since the latter uses mitochondrial metabolism as a marker of cell viability, and we sought to avoid any potential confounding effects of culture conditions on mitochondria.

### In vivo studies

GBM12, GBM164, and GBM196 cells were sourced from tumors propagated as subcutaneous growths in athymic nude mice and prepared for implantation. Briefly, after euthanizing each animal, tumors were aseptically excised from the flank and minced in a sterile culture dish with a scalpel. The cell suspension was centrifuged after mechanical disruption, filtered through 70 µM nylon-mesh filters, re-centrifuged, and re-suspended in an equal volume ratio of cell culture media to Matrigel. A 16-gauge needle and syringe were used to inject the cell suspension into the flanks of 6 week-old female NCr athymic nude mice (NCRNU-F sp/sp, Taconic).

Intracranial injections of NPA and NPAC1 cells were performed as described previously in 10 week-old female C57BL/6J mice (Jackson) [17].

Animals received post-operative support care through administration of 0.9% saline solution, DietGel 76A, and thermal support while recovering from anesthesia until awake and ambulatory. One administration of meloxicam 1 mg/kg was given at the time of the procedure for 24 hour analgesia coverage, followed by another dose approximately 24 hours post-procedure, and a final dose 48 hours post-procedure if deemed necessary.

### Animal monitoring and treatment

All animals were fed standard rodent chow (standard diet, or SD) for 3 days following flank or intracerebral engraftment before being randomized to either remain on SD *ad libitum*, or given a formulated KD (Ketogenic Diet TD.96355, Envigo Teklad Diets, Madison WI) *ad libitum*. The KD was received directly from the manufacturer, and was a nutritionally complete diet composed of 15.3% protein, 0.5% carbohydrate, and 67.4% fat by weight (**Table 2**). Mice assigned to the KD group were kept on KD for one week, followed by SD for one week, and so on, in order to prevent obesity and reduce midlife mortality. Mice were still allowed to feed *ad libitum*, in order to reduce potentially confounding effects of weight loss from calorie-restricted diets and more effectively maintain plasma ketone levels, as was described by others [13, 18]. At the end of each week, blood from the saphenous vein was collected to test circulating metabolite levels using the Precision Xtra Blood Glucose and Ketone Monitoring System (Abbott SKU# 9881465). Animals were monitored twice weekly for signs of morbidity, such as weight loss, behavioral changes, and hunched positions, and were euthanized when tumor size reached 2,000 mm^3^ (below the maximum IACUC-approved volume), or when moribund. Euthanasia was performed via CO_2_ asphyxiation followed by cervical dislocation, in accordance with IACUC-approved guidelines.

**Table 2:** Summary of macronutrient composition of standard and experimental diets presented as proportion of total kilocalories.

| Diets | Standard Diet | Ketogenic Diet |
| --- | --- | --- |
| Macronutrients | Envigo Teklad LM-485 Rodent Diet | Envigo Teklad Ketogenic Diet td.96355 |
| Protein % | 19.1 | 9.2 |
| Carbohydrate % | 44.3 | 0.3 |
| Fat % | 5.8 | 90.5 |
| kcal/g | 3.1 | 6.7 |

Flank tumor volume was based on caliper measurements and calculated using the modified ellipsoidal formula [19, 20]:

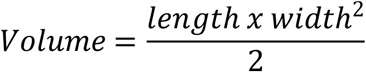

### Statistical Analyses

Differences between mean values of two groups were compared using unpaired two-sample *t*-tests, or for interactions between glucose and ketones *in vitro*, via two-way ANOVA; *P* values less than 0.05 were considered significant. Log-rank tests compared survival between groups. Graph generation and statistical analyses were performed with GraphPad Prism 9 (GraphPad Software, San Diego, CA).

## RESULTS

First, we examined the effects of a ketogenic-like diet on cultured IDH1^wt^ and IDH1^mut^ patient-derived cancer cell lines under the following conditions: (i) normal basal glucose (NG) of 4.5 g/L; (ii) low glucose (LG) of 1.0 g/L; (iii) NG with 10 mM β-hydroxybutyrate (BHB); (iv) LG with BHB (**Figure 1**). Both pairs of IDH1^wt^ and IDH1^mut^ cell types responded similarly, with slightly attenuated proliferation in LG medium, but no effect of BHB in either NG or LG medium, as indicated by two-way ANOVA analyses (**Table 3**). Specifically, among IDH1^wt^ GBM6 cells, proliferation was higher in NG than LG (*F* (1, 8)=10.0, *P*=0.013), but BHB had no effect in either NG or LG media (*F* (1,8)=0.20, *P*=0.66). There was also no statistically significant interaction between glucose and BHB (*F* (1,8)=0.0048, *P*=0.95). Among IDH1^wt^ GBM43 cells, proliferation was higher in NG than LG medium (*F* (1, 8)=12.0, *P*=0.0088), but BHB had no effect in either NG or LG media (*F* (1,8)=0.48, *P*=0.51), with no significant interaction between glucose and BHB (*F* (1,8)=0.044, *P*=0.84). Among IDH1^mut^ TB09 cells, proliferation was higher in NG than LG (*F* (1, 8)=42.0, *P*=0.0002), but BHB had no effect in either NG or LG media (*F* (1,8)=1.4, *P*=0.28), and there was no statistically significant interaction between glucose and BHB (*F* (1,8)=0.90, *P*=0.37). Similarly, IDH1^mut^ HT1080 cells showed higher proliferation in NG than LG medium (*F* (1, 8)=13.0, *P*=0.0069), but BHB had no effect in either NG or LG media (*F* (1,8)=2.2, *P*=0.18). There was also no significant interaction between glucose and BHB (*F* (1,8)=1.8, *P*=0.22).

**Figure 1.**
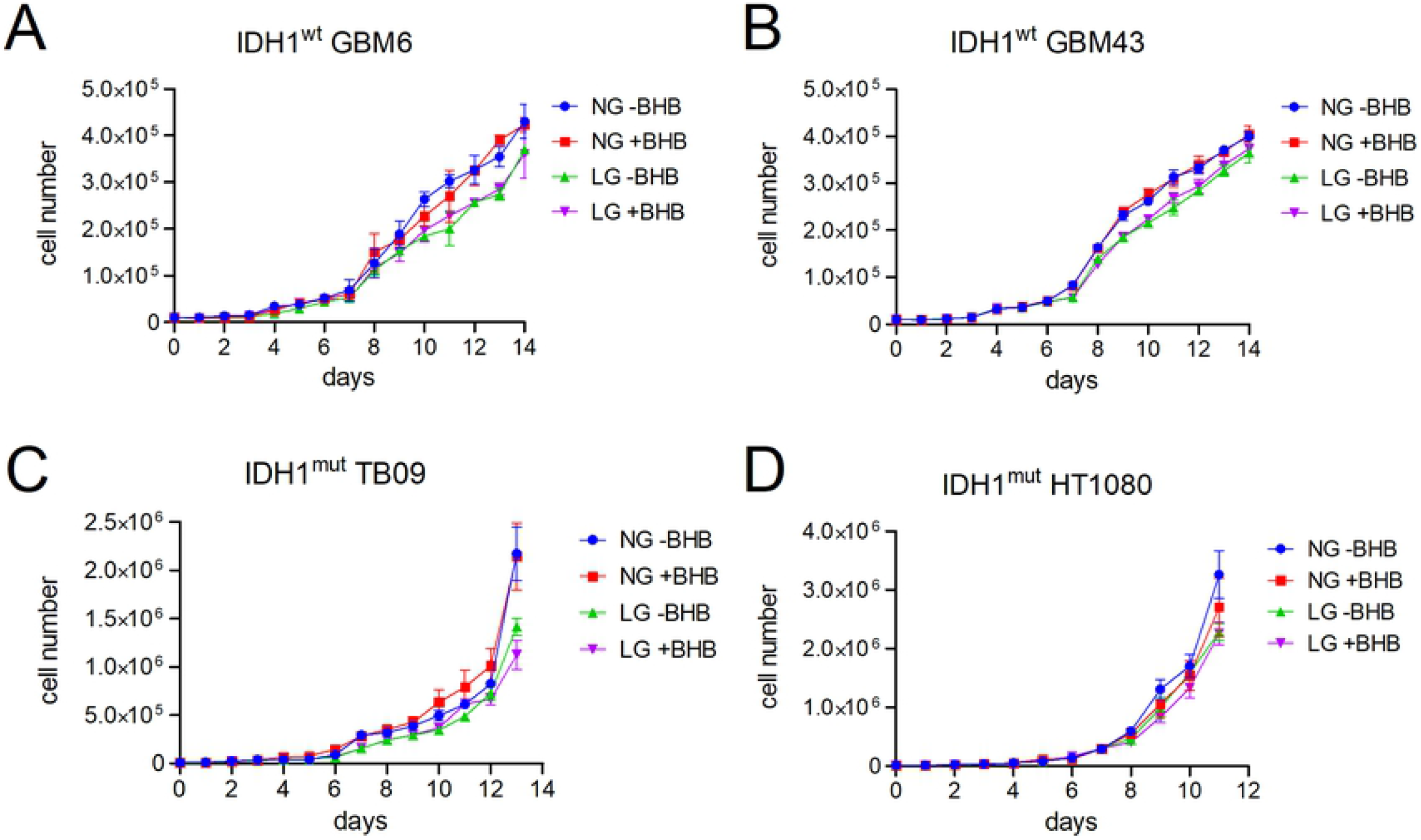
Effect of ketones on patient-derived IDH1^wt^ and IDH1^mut^ tumor cell proliferation *in vitro*. Absolute cell counts over 14 days of culture in either normal glucose (NG) or low glucose (LG), with or without 10 mM β-hydroxybutyrate (BHB). Each data point is shown as mean ±SEM. Two-way ANOVA analyses of the final time points are described in the Results text and Table 3.

**Table 3:** Two-way ANOVA analyses of *in vitro* proliferation of IDH^wt^ and IDH^mut^ tumor cells in varying glucose and ketogenic culture conditions. NG=4.5 g/L, LG=1.0 g/L, -BHB=without β-hydroxybutyrate, +BHB=10 mM β-hydroxybutyrate. Each analysis was done on the last day of culture for each cell type,as indicated in Figure 1.

| cell type | source of variation | % of total variation | *P* |
| --- | --- | --- | --- |
| **IDH<sup>wt</sup> GBM6** | interaction | 0.026 | 0.95 |
|  | *NG* vs. *LG* | 55.0 | 0.013 |
|  | -BHB vs. +BHB | 1.1 | 0.66 |
| **IDH<sup>wt</sup> GBM43** | interaction | 0.21 | 0.84 |
|  | *NG* vs. *LG* | 58.0 | 0.0088 |
|  | -BHB vs. +BHB | 2.4 | 0.51 |
| **IDH<sup>mut</sup> TB09** | interaction | 1.7 | 0.37 |
|  | *NG* vs. *LG* | 80.0 | 0.0002 |
|  | -BHB vs. +BHB | 2.6 | 0.28 |
| **IDH<sup>mut</sup> HT1080** | interaction | 7.2 | 0.22 |
|  | *NG* vs. *LG* | 52.0 | 0.0069 |
|  | -BHB vs. +BHB | 8.7 | 0.18 |

Next, we evaluated the ability of a cycling KD (see “Methods”) to induce ketosis in mice without tumors. Over the first 5 weeks that mice were on KD, blood glucose levels were similar to control mice on SD, even during weeks in which KD was implemented (**Figure 2A**). Interestingly, blood glucose rose 15% in KD mice after week 6 (147.8 mg/dl in KD versus 128.0 mg/dl in SD, *P*=0.0029) (**Figure 2A**). This persisted in week 7 (144.3 mg/dl in KD versus 127.3 mg/dl in SD, *P*=0.0053). Ketones, in contrast, increased by 112.5% after just one week on KD compared to mice on SD (1.13 mM versus 0.53 mM, *P*=0.00022) (**Figure 2B**). While circulating ketones declined in KD mice during the weeks when they were back on SD, they remained 81-236% higher than in SD mice over the entire 7-week interval, consistent with previously published data by other who employed this type of KD [13, 18]. Body mass was 13% higher in KD mice than SD mice by week 4 (26.3 grams versus 23.3 grams, *P*=0.033), but by week 6, the mass of the SD mice matched the KD mice (**Figure 2C**).

**Figure 2:**
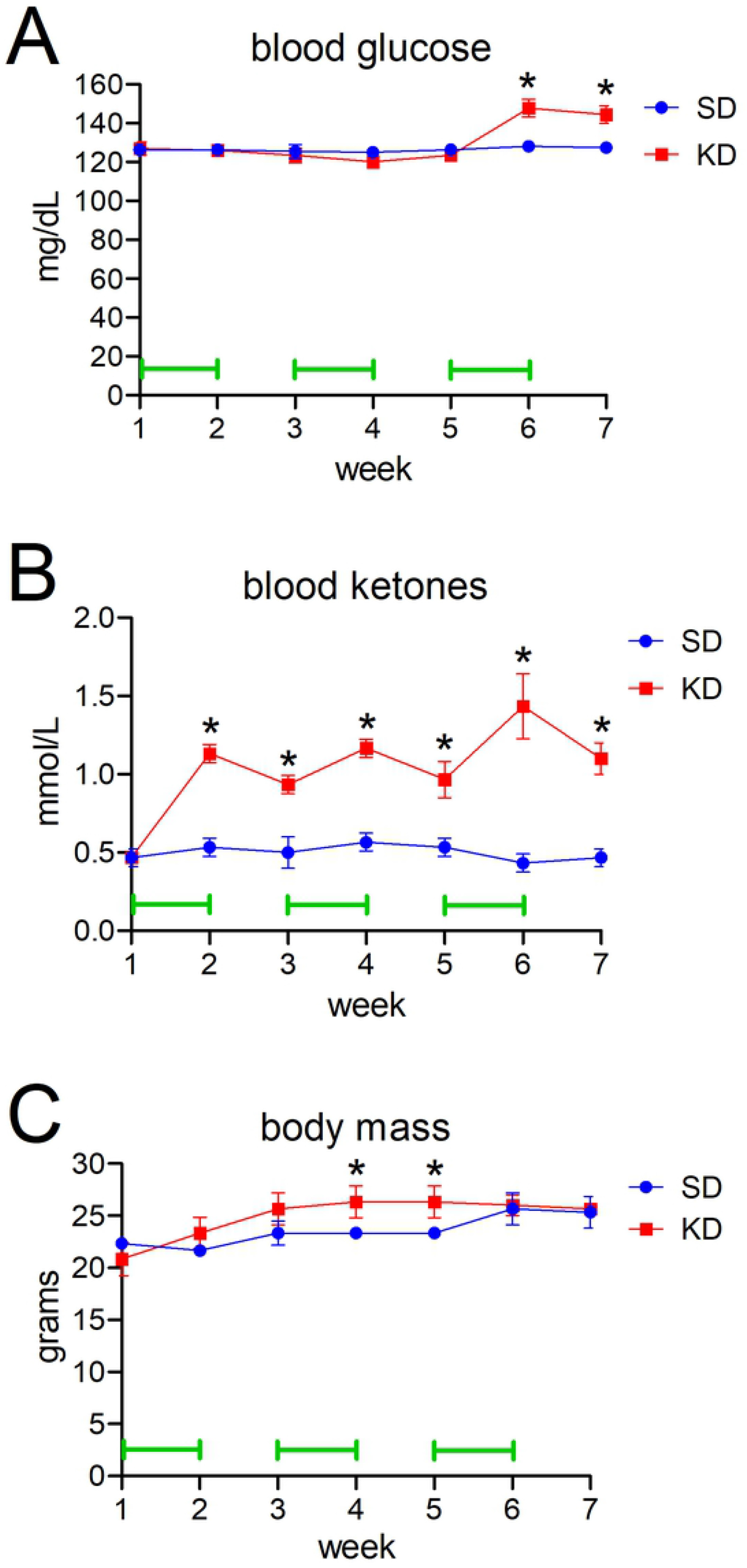
Basic metabolic parameters of mice on a cycling KD. Saphenous vein blood samples tested for glucose (A) and ketones (B) at the end of each week. Mouse body weights are shown in (C). Green bars indicate weeks in which mice were on KD.

Mice bearing subcutaneous flank patient-derived xenografts (PDX) of endogenous IDH1^wt^ GBM12, or IDH1^mut^ grade 4 astrocytoma (GBM164 and GBM196), were subjected to a cycling KD or kept on regular SD. (The flank was chosen because IDH1^mut^ GBM164 and GBM196 cells grow poorly intracerebrally, and these two PDX models were chosen for *in vivo* studies because IDH1^mut^ TB09 cells grow poorly *in vivo*.) As expected, IDH1^wt^ GBM12 grew faster than IDH^mut^ GBM164 and GBM196 (**Figure 3**). However, within each PDX subtype, growth was similar in KD mice and SD mice (GBM164: 221 mm^3^ SD versus 226 mm^3^ KD, *P*=0.95; GBM196: 354 mm^3^ SD versus 335 mm^3^ KD, *P*=0.90; GBM12: 1540 mm^3^ SD versus 1251 mm^3^ KD, *P*=0.44).

**Figure 3.**
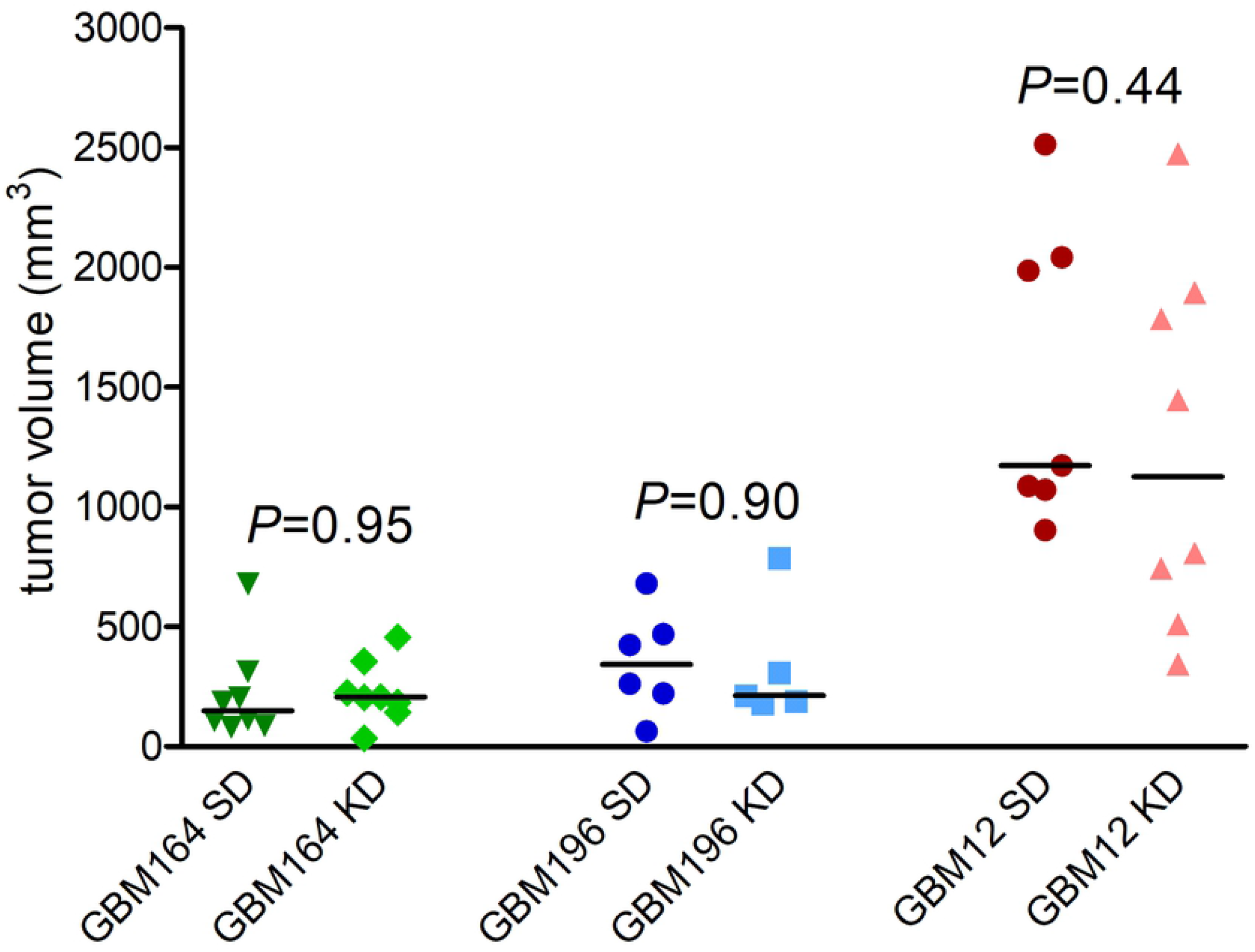
The effect of cycling KD on patient-derived IDH1^wt^ and IDH1^mut^ flank xenograft growth. Tumor volumes for mice subcutaneously engrafted with IDH1^mut^ GBM164 at 21 days, IDH1^mut^ GBM 196 at 14 days, and IDH1^wt^ GBM12 at 14 days, either on SD or cycling KD. Time points differed due to differing rates of tumor growth. Bars=means, *P* values calculated by unpaired t-tests.

To determine the effect of KD on intracerebral, isogenic-matched IDH1^wt^ and IDH1^mut^ gliomas, we engrafted IDH1^wt^ NPA and IDH1^mut^ NPAC1 cells (**Table 1**) into the brains of immunocompetent mice (**Figure 4**). Among mice maintained on SD, those engrafted with IDH1^mut^ NPAC1 cells survived 20% longer than mice engrafted with IDH1^wt^ NPA cells (median survival 26.5 days versus 22.0 days, HR=0.11, 95% CI=0.03-0.39, *P*=0.0022), in keeping with published data on these models [17]. However, KD had no effect on survival in either NPA-engrafted subjects (HR=0.47, 95% CI=0.15-1.5, *P*=0.27) (**Figure 4A**) or NPAC1-engrafted subjects (HR=1.0, 95% CI=0.31-3.2, *P*=0.81) (**Figure 4B**).

**Figure 4.**
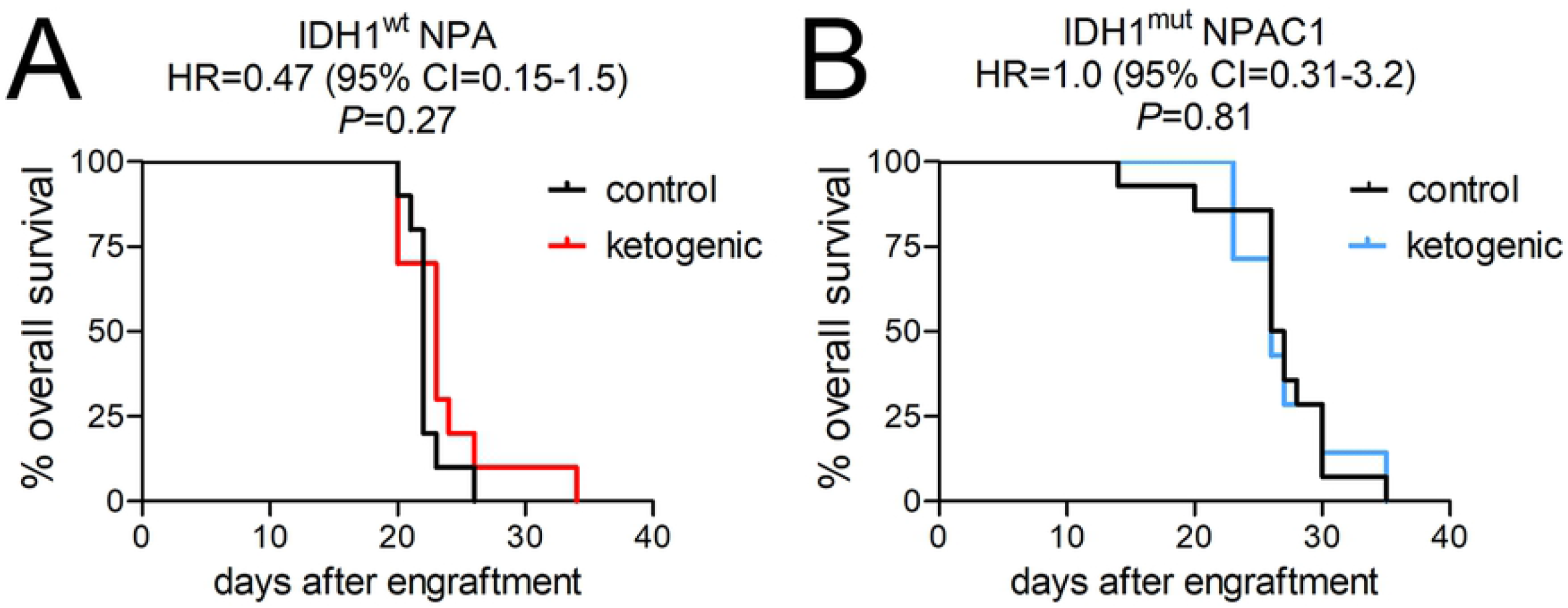
The effect of cycling KD on patient-derived IDH1^wt^ and IDH1^mut^ intracranial xenograft growth. Kaplan-Meier survival curves of mice engrafted with (A) IDH1^wt^ NPA or (B) IDH1^mut^ NPAC1, either on SD or cycling KD.

## DISCUSSION

Altered metabolism in cancer cells raises the possibility of exploiting metabolic vulnerabilities to inhibit tumor growth, such as putting patients on KD. However, while KD may have efficacy against some cancers, clinical studies have been limited in glioma patients, and have so far mostly focused on feasibility, not outcomes [15, 21-26]. At the preclinical level, some studies showed that mice engrafted with IDH1^wt^ GBM did not benefit from KD [13, 27], although others have suggested otherwise [28], and to the best of our knowledge, there has not yet been a direct experimental comparison of IDH1^wt^ and IDH1^mut^ gliomas exposed to KD. Another study by others suggested that IDH1^mut^ might confer sensitivity to KD, based on *in vitro* data using HCT116 cells engineered to express IDH1 R132H [14]. Thus, we sought to determine KD efficacy in a variety of preclinical *in vitro* and *in vivo* models of IDH1^wt^ and IDH1^mut^ glioma. Our data suggest that KD does not have significant activity against either IDH^wt^ or IDH^mut^ glioma.

Since the original studies describing IDH1^mut^ and its neoenzymatic activity in cancer, a great deal of research has been done studying the metabolic effects of IDH1^mut^, often generating conflicting results. For example, while some have shown that IDH1^mut^ depletes the cell of TCA intermediates [29-34], others have found little to no changes in those intermediates [10, 35, 36]; indeed, IDH1^mut^ gliomas may actually use lactate and glutamate anaplerosis to replenish TCA intermediates [35]. Some have suggested that glycolysis is reduced in IDH^mut^ gliomas [34, 36], but one group found reduced glucose uptake by IDH^mut^ glioma cells, and otherwise no difference in the rate of glycolysis compared to IDH^wt^ cells [37].

Results in IDH^mut^ metabolic research seem to vary greatly depending on whether one is studying cells artificially overexpressing IDH^mut^, or is focusing on cells and tissues with endogenous IDH^mut^, as more pronounced metabolic changes tend to occur in the former than the latter [37]. This suggests that cells with naturally-occurring IDH^mut^ can, over time, adjust their metabolism to at least partially compensate for perturbations caused by the mutant enzyme. For example, IDH^mut^ gliomas may adjust for the depletion of TCA metabolites by upregulating glutamate dehydrogenase 2 expression [34]. Since IDH^mut^ gliomas mostly use the TCA precursor glutamine to produce D2HG, these tumors compensate by turning pyruvate into TCA chemicals [38, 39]. Although IDH^mut^ consumes NAPDH, which should lead to glutathione depletion, IDH^mut^ gliomas upregulate enzymes involved in glutathione synthesis, thereby maintaining glutathione levels [40]. Thus, when IDH^wt^ wild-type cells are abruptly forced to overexpress IDH^mut^, any metabolic results, including sensitivity to KD-like conditions, need to be validated in patient-derived and/or transgenic models with endogenous IDH^mut^.

Strict adherence to KD is notoriously difficult [15, 41]. Most prescribed KD includes caloric restriction, which can lead to undesired weight loss and introduce confounding effects on metabolism. Even adjusting dietary supplementation based on measured body weight has been shown to significantly affect overall body weight [13]. A recent patient-based study showed that KD does cause similarly elevated ketones within IDH^wt^ and IDH^mut^ gliomas by magnetic resonance spectroscopy [15], but patient survival was not a part of that analysis. Other recent preclinical and clinical studies have suggested that gliomas can metabolically adapt to ketogenesis, and even use ketones to facilitate disease progression [25, 27]. Our experimental results align with such studies, and show that, despite achieving a ketosis-like state in tumor bearing animals, no difference in overall tumor volume or survival benefit was observed. Thus, our data suggest that KD may not have significant effects on the outcomes of glioma patients, regardless of IDH^mut^ status.

## ACKNOWLEGMENTS

This work was supported by R01NS102669 (CH), R01NS117104 (CH), R01NS118039 (CH), the Northwestern University P50CA221747 SPORE in Brain Tumor Research, and the Lou and Jean Malnati Brain Tumor Institute.

## REFERENCES

1. Ostrom QT, Cioffi G, Gittleman H, Patil N, Waite K, Kruchko C, et al. CBTRUS Statistical Report: Primary Brain and Other Central Nervous System Tumors Diagnosed in the United States in 2012-2016. Neuro Oncol. 2019;21(Suppl 5):v1–v100. Epub 2019/11/02. doi: 10.1093/neuonc/noz150. PubMed PMID: 31675094; PubMed Central PMCID: PMCPMC6823730.

2. Stupp R, Mason WP, van den Bent MJ, Weller M, Fisher B, Taphoorn MJ, et al. Radiotherapy plus concomitant and adjuvant temozolomide for glioblastoma. N Engl J Med. 2005;352(10):987–96. PubMed PMID: 15758009.

3. Unruh D, Mirkov S, Wray B, Drumm M, Lamano J, Li YD, et al. Methylation-dependent Tissue Factor suppression contributes to the reduced malignancy of IDH1 mutant gliomas. Clin Cancer Res. 2018;28:1078–0432.

4. Burnet NG, Jefferies SJ, Benson RJ, Hunt DP, Treasure FP. Years of life lost (YLL) from cancer is an important measure of population burden--and should be considered when allocating research funds. Br J Cancer. 2005;92(2):241–5.

5. Hanahan D, Weinberg RA. Hallmarks of cancer: the next generation. Cell. 2011;144(5):646–74. PubMed PMID: 21376230.

6. Warburg O, Wind F, Negelein E. THE METABOLISM OF TUMORS IN THE BODY. J Gen Physiol. 1927;8(6):519–30. PubMed PMID: 19872213.

7. Horbinski C. What do we know about IDH1/2 mutations so far, and how do we use it? Acta Neuropathol. 2013;125(5):621–36. Epub 2013/03/21. doi: 10.1007/s00401-013-1106-9 [doi]. PubMed PMID: 23512379.

8. Parsons DW, Jones S, Zhang X, Lin JC, Leary RJ, Angenendt P, et al. An integrated genomic analysis of human glioblastoma multiforme. Science. 2008;321(5897):1807–12. PubMed PMID: 18772396.

9. Yan H, Parsons DW, Jin G, McLendon R, Rasheed BA, Yuan W, et al. IDH1 and IDH2 mutations in gliomas. N Engl J Med. 2009;360(8):765–73. PubMed PMID: 19228619.

10. Dang L, White DW, Gross S, Bennett BD, Bittinger MA, Driggers EM, et al. Cancer-associated IDH1 mutations produce 2-hydroxyglutarate. Nature. 2009;462(7274):739–44. PubMed PMID: 19935646.

11. Gano LB, Patel M, Rho JM. Ketogenic diets, mitochondria, and neurological diseases. J Lipid Res. 2014;55(11):2211–28. Epub 2014/05/23. doi: 10.1194/jlr.R048975. PubMed PMID: 24847102; PubMed Central PMCID: PMCPMC4617125.

12. Chung HY, Park YK. Rationale, Feasibility and Acceptability of Ketogenic Diet for Cancer Treatment. J Cancer Prev. 2017;22(3):127–34. Epub 2017/10/12. doi: 10.15430/jcp.2017.22.3.127. PubMed PMID: 29018777; PubMed Central PMCID: PMCPMC5624453.

13. De Feyter HM, Behar KL, Rao JU, Madden-Hennessey K, Ip KL, Hyder F, et al. A ketogenic diet increases transport and oxidation of ketone bodies in RG2 and 9L gliomas without affecting tumor growth. Neuro Oncol. 2016;18(8):1079–87. Epub 2016/05/05. doi: 10.1093/neuonc/now088. PubMed PMID: 27142056; PubMed Central PMCID: PMCPMC4933488.

14. Fu X, Chin RM, Vergnes L, Hwang H, Deng G, Xing Y, et al. 2-Hydroxyglutarate Inhibits ATP Synthase and mTOR Signaling. Cell Metab. 2015;22(3):508–15. Epub 2015/07/21. doi: 10.1016/j.cmet.2015.06.009. PubMed PMID: 26190651; PubMed Central PMCID: PMCPMC4663076.

15. Schreck KC, Hsu FC, Berrington A, Henry-Barron B, Vizthum D, Blair L, et al. Feasibility and Biological Activity of a Ketogenic/Intermittent-Fasting Diet in Patients With Glioma. Neurology. 2021. Epub 2021/07/09. doi: 10.1212/wnl.0000000000012386. PubMed PMID: 34233941.

16. Vaubel RA, Tian S, Remonde D, Schroeder MA, Mladek AC, Kitange GJ, et al. Genomic and Phenotypic Characterization of a Broad Panel of Patient-Derived Xenografts Reflects the Diversity of Glioblastoma. Clin Cancer Res. 2020;26(5):1094–104. Epub 2019/12/20. doi: 10.1158/1078-0432.Ccr-19-0909. PubMed PMID: 31852831; PubMed Central PMCID: PMCPMC7056576.

17. Nunez FJ, Mendez FM, Kadiyala P, Alghamri MS, Savelieff MG, Garcia-Fabiani MB, et al. IDH1-R132H acts as a tumor suppressor in glioma via epigenetic up-regulation of the DNA damage response. Sci Transl Med. 2019;11(479). Epub 2019/02/15. doi: 10.1126/scitranslmed.aaq1427. PubMed PMID: 30760578; PubMed Central PMCID: PMCPMC6400220.

18. Newman JC, Covarrubias AJ, Zhao M, Yu X, Gut P, Ng CP, et al. Ketogenic Diet Reduces Midlife Mortality and Improves Memory in Aging Mice. Cell Metab. 2017;26(3):547–57 e8. PubMed PMID: 28877458.

19. Faustino-Rocha A, Oliveira PA, Pinho-Oliveira J, Teixeira-Guedes C, Soares-Maia R, da Costa RG, et al. Estimation of rat mammary tumor volume using caliper and ultrasonography measurements. Lab Anim (NY). 2013;42(6):217–24. Epub 2013/05/22. doi: 10.1038/laban.254. PubMed PMID: 23689461.

20. Jensen MM, Jørgensen JT, Binderup T, Kjaer A. Tumor volume in subcutaneous mouse xenografts measured by microCT is more accurate and reproducible than determined by 18F-FDG-microPET or external caliper. BMC Med Imaging. 2008;8:16. Epub 2008/10/18. doi: 10.1186/1471-2342-8-16. PubMed PMID: 18925932; PubMed Central PMCID: PMCPMC2575188.

21. Klement RJ, Brehm N, Sweeney RA. Ketogenic diets in medical oncology: a systematic review with focus on clinical outcomes. Med Oncol. 2020;37(2):14. Epub 2020/01/14. doi: 10.1007/s12032-020-1337-2. PubMed PMID: 31927631.

22. Foppiani A, De Amicis R, Lessa C, Leone A, Ravella S, Ciusani E, et al. Isocaloric Ketogenic Diet in Adults with High-Grade Gliomas: A Prospective Metabolic Study. Nutr Cancer. 2021;73(6):1004–14. Epub 2021/03/11. doi: 10.1080/01635581.2020.1779759. PubMed PMID: 33689522.

23. Porper K, Shpatz Y, Plotkin L, Pechthold RG, Talianski A, Champ CE, et al. A Phase I clinical trial of dose-escalated metabolic therapy combined with concomitant radiation therapy in high-grade glioma. Journal of neuro-oncology. 2021;153(3):487–96. Epub 2021/06/22. doi: 10.1007/s11060-021-03786-8. PubMed PMID: 34152528.

24. Perez A, van der Louw E, Nathan J, El-Ayadi M, Golay H, Korff C, et al. Ketogenic diet treatment in diffuse intrinsic pontine glioma in children: Retrospective analysis of feasibility, safety, and survival data. Cancer Rep (Hoboken). 2021:e1383. Epub 2021/05/04. doi: 10.1002/cnr2.1383. PubMed PMID: 33939330.

25. Wenger KJ, Wagner M, Harter PN, Franz K, Bojunga J, Fokas E, et al. Maintenance of Energy Homeostasis during Calorically Restricted Ketogenic Diet and Fasting-MR-Spectroscopic Insights from the ERGO2 Trial. Cancers (Basel). 2020;12(12). PubMed PMID: 33261052.

26. Martin-McGill KJ, Marson AG, Tudur Smith C, Young B, Mills SJ, Cherry MG, et al. Ketogenic diets as an adjuvant therapy for glioblastoma (KEATING): a randomized, mixed methods, feasibility study. Journal of neuro-oncology. 2020;147(1):213–27. PubMed PMID: 32036576.

27. Sperry J, Condro MC, Guo L, Braas D, Vanderveer-Harris N, Kim KKO, et al. Glioblastoma Utilizes Fatty Acids and Ketone Bodies for Growth Allowing Progression during Ketogenic Diet Therapy. iScience. 2020;23(9):101453. PubMed PMID: 32861192.

28. Ciusani E, Vasco C, Rizzo A, Girgenti V, Padelli F, Pellegatta S, et al. MR-Spectroscopy and Survival in Mice with High Grade Glioma Undergoing Unrestricted Ketogenic Diet. Nutr Cancer. 2020:1–8. Epub 2020/09/22. doi: 10.1080/01635581.2020.1822423. PubMed PMID: 32954880.

29. Biedermann J, Preussler M, Conde M, Peitzsch M, Richter S, Wiedemuth R, et al. Mutant IDH1 Differently Affects Redox State and Metabolism in Glial Cells of Normal and Tumor Origin. Cancers (Basel). 2019;11(12). Epub 2020/01/01. doi: 10.3390/cancers11122028. PubMed PMID: 31888244; PubMed Central PMCID: PMCPMC6966450.

30. Grassian AR, Parker SJ, Davidson SM, Divakaruni AS, Green CR, Zhang X, et al. IDH1 mutations alter citric acid cycle metabolism and increase dependence on oxidative mitochondrial metabolism. Cancer Res. 2014;74(12):3317–31. Epub 2014/04/24. doi: 10.1158/0008-5472.Can-14-0772-t. PubMed PMID: 24755473; PubMed Central PMCID: PMCPMC4885639.

31. Miyata S, Tominaga K, Sakashita E, Urabe M, Onuki Y, Gomi A, et al. Comprehensive Metabolomic Analysis of IDH1(R132H) Clinical Glioma Samples Reveals Suppression of β-oxidation Due to Carnitine Deficiency. Sci Rep. 2019;9(1):9787. Epub 2019/07/07. doi: 10.1038/s41598-019-46217-5. PubMed PMID: 31278288; PubMed Central PMCID: PMCPMC6611790.

32. Ohka F, Ito M, Ranjit M, Senga T, Motomura A, Motomura K, et al. Quantitative metabolome analysis profiles activation of glutaminolysis in glioma with IDH1 mutation. Tumour Biol. 2014;35(6):5911–20. Epub 2014/03/05. doi: 10.1007/s13277-014-1784-5. PubMed PMID: 24590270.

33. Reitman ZJ, Jin G, Karoly ED, Spasojevic I, Yang J, Kinzler KW, et al. Profiling the effects of isocitrate dehydrogenase 1 and 2 mutations on the cellular metabolome. Proc Natl Acad Sci U S A. 2011;108(8):3270–5. PubMed PMID: 21289278.

34. Waitkus MS, Pirozzi CJ, Moure CJ, Diplas BH, Hansen LJ, Carpenter AB, et al. Adaptive Evolution of the GDH2 Allosteric Domain Promotes Gliomagenesis by Resolving IDH1(R132H)-Induced Metabolic Liabilities. Cancer Res. 2018;78(1):36–50. Epub 2017/11/04. doi: 10.1158/0008-5472.Can-17-1352. PubMed PMID: 29097607; PubMed Central PMCID: PMCPMC5754242.

35. Khurshed M, Molenaar RJ, Lenting K, Leenders WP, van Noorden CJF. In silico gene expression analysis reveals glycolysis and acetate anaplerosis in IDH1 wild-type glioma and lactate and glutamate anaplerosis in IDH1-mutated glioma. Oncotarget. 2017;8(30):49165–77. Epub 2017/05/04. doi: 10.18632/oncotarget.17106. PubMed PMID: 28467784; PubMed Central PMCID: PMCPMC5564758.

36. Zhou L, Wang Z, Hu C, Zhang C, Kovatcheva-Datchary P, Yu D, et al. Integrated Metabolomics and Lipidomics Analyses Reveal Metabolic Reprogramming in Human Glioma with IDH1 Mutation. J Proteome Res. 2019;18(3):960–9. Epub 2019/01/01. doi: 10.1021/acs.jproteome.8b00663. PubMed PMID: 30596429.

37. Garrett M, Sperry J, Braas D, Yan W, Le TM, Mottahedeh J, et al. Metabolic characterization of isocitrate dehydrogenase (IDH) mutant and IDH wildtype gliomaspheres uncovers cell type-specific vulnerabilities. Cancer Metab. 2018;6:4. Epub 2018/04/26. doi: 10.1186/s40170-018-0177-4. PubMed PMID: 29692895; PubMed Central PMCID: PMCPMC5905129.

38. Izquierdo-Garcia JL, Cai LM, Chaumeil MM, Eriksson P, Robinson AE, Pieper RO, et al. Glioma cells with the IDH1 mutation modulate metabolic fractional flux through pyruvate carboxylase. PLoS One. 2014;9(9):e108289. Epub 2014/09/23. doi: 10.1371/journal.pone.0108289. PubMed PMID: 25243911; PubMed Central PMCID: PMCPMC4171511.

39. Izquierdo-Garcia JL, Viswanath P, Eriksson P, Cai L, Radoul M, Chaumeil MM, et al. IDH1 Mutation Induces Reprogramming of Pyruvate Metabolism. Cancer Res. 2015;75(15):2999–3009. Epub 2015/06/06. doi: 10.1158/0008-5472.Can-15-0840. PubMed PMID: 26045167; PubMed Central PMCID: PMCPMC4526330.

40. Fack F, Tardito S, Hochart G, Oudin A, Zheng L, Fritah S, et al. Altered metabolic landscape in IDH-mutant gliomas affects phospholipid, energy, and oxidative stress pathways. EMBO Mol Med. 2017;9(12):1681–95. Epub 2017/10/22. doi: 10.15252/emmm.201707729. PubMed PMID: 29054837; PubMed Central PMCID: PMCPMC5709746.

41. Brouns F. Overweight and diabetes prevention: is a low-carbohydrate-high-fat diet recommendable? Eur J Nutr. 2018;57(4):1301–12. PubMed PMID: 29541907.

